# Prior cocaine use diminishes encoding of latent information by orbitofrontal, but not medial, prefrontal ensembles

**DOI:** 10.1101/2024.02.21.581248

**Authors:** Lauren E. Mueller, Caitlin Konya, Melissa J. Sharpe, Andrew M. Wikenheiser, Geoffrey Schoenbaum

## Abstract

Maladaptive decision-making is a hallmark of substance use disorders, though how drugs of abuse alter neural representations supporting adaptive behavior remains poorly understood. Past studies show the orbitofrontal (OFC) and prelimbic (PL) cortices are important for decision making, tracking both task-relevant and latent information. However, previous studies have focused on how drugs of abuse impact the firing rates of individual units. More work at the ensemble level is necessary to accurately characterize potential drug-induced changes. Using single-unit recordings in rats during a multidimensional decision-making task and then applying population and ensemble level analyses, we show that prior use of cocaine altered the strength and structure of task-relevant and latent representations in the OFC, changes relatable to suboptimal decision making in this and perhaps other settings. These data expand our understanding of the neuropathological underpinnings of maladaptive decision-making in SUDs, potentially enabling enhanced future treatment strategies.

## Introduction

Maladaptive decision making is a prominent component of substance use disorders (SUDs), demonstrated by continued drug use, pathological gambling, and other risky behaviors in the face of adverse consequences ^1–3^. Previous work has linked these symptoms to effects of prior drug experience on choice in reversal tasks in humans ^4, 5^, rats ^6, 7^, mice ^8^, and monkeys ^9, 10^. Rats with prior drug histories also demonstrate impairments in settings such as outcome devaluation^11, 12^, contingency degradation ^13, 14^, sensory preconditioning ^15^, and Pavlovian-to-instrumental transfer ^16^. These impairments may explain why animals and humans behave suboptimally across a wide range of scenarios following drug experience.

However, the neural underpinnings of these impairments are not well-understood. The orbitofrontal and prelimbic cortices (OFC and PL) are prefrontal regions vital for normal decision making. Further, the OFC and PL play important roles in tracking not only task-relevant associative information but also information that is hidden (i.e. not directly observable) or latent (i.e. not necessary at the time of learning for task performance). The special role these areas play in protecting such information is evident in their necessity for normal performance in a variety of settings, such as devaluation^17–19^, sensory preconditioning^20, 21^, over-expectation^22, 23^, contingency degradation^24–27^, latent inhibition^28–30^, and transfer^31–33^. Drugs of abuse persistently impair the ability of these regions to process associative information in these settings ^34–36^, effects evident at both the behavioral ^6, 12, 34, 37–39^ and neurophysiological levels ^40–43^. However, existing studies have focused on characterizing individual unit firing rate changes, measures which may not be nuanced enough to paint an accurate picture of drug-induced informational processing deficits. Larger population or ensemble-level analyses, which fully describe the type and structure of information being represented – including latent information - are necessary.

Here we addressed these questions by recording single-unit activity in a multidimensional decision-making task and applying ensemble analyses to test the strength and structure of the representation of both task-relevant and latent information. We found that prior cocaine experience significantly altered both, particularly within the OFC.

## Results

Rats were shaped on an odor-guided decision-making task in which rewards delivered at two wells changed across blocks of trials (Fig. 1a) ^44^. The paradigm consisted of a short warm-up block, followed by four full-length trial blocks across a single session. On each trial, one of three odor cues was presented, and rats were required to respond to the left or to the right fluid well on forced-choice trials, or in either direction on free-choice trials, to receive milk rewards that differed in size (big: three 0.05 ml drops; small: one 0.05 ml drop) and flavor (vanilla or chocolate milk). At unsignaled block switches, the reward available in each fluid well changed in either size (i.e., value block switch) or flavor (i.e., identity block switch), establishing distinct sets of response-outcome contingencies per trial block. After acquiring this task, rats were trained to self-administer either oral sucrose (n = 4) or cocaine (n = 4) on a fixed ratio one schedule of reinforcement for 3 h/d for 14 d (Supplemental Figs. 1 and 2). Following self-administration, rats received some additional training on the decision-making paradigm, after which two drivable bundles of electrodes were implanted into each rat, one in the OFC and one in the PL, with the hemisphere of the implanted regions counterbalanced across rats (i.e., left OFC, right PL or left PL, right OFC; Supplemental Fig. 1). After recovery, rats were given seven days of reminder training for acclimation to recording cables and then neural recording sessions began (Supplemental Fig. 1); electrodes were advanced after each session to record new neurons.

**Figure 1.**
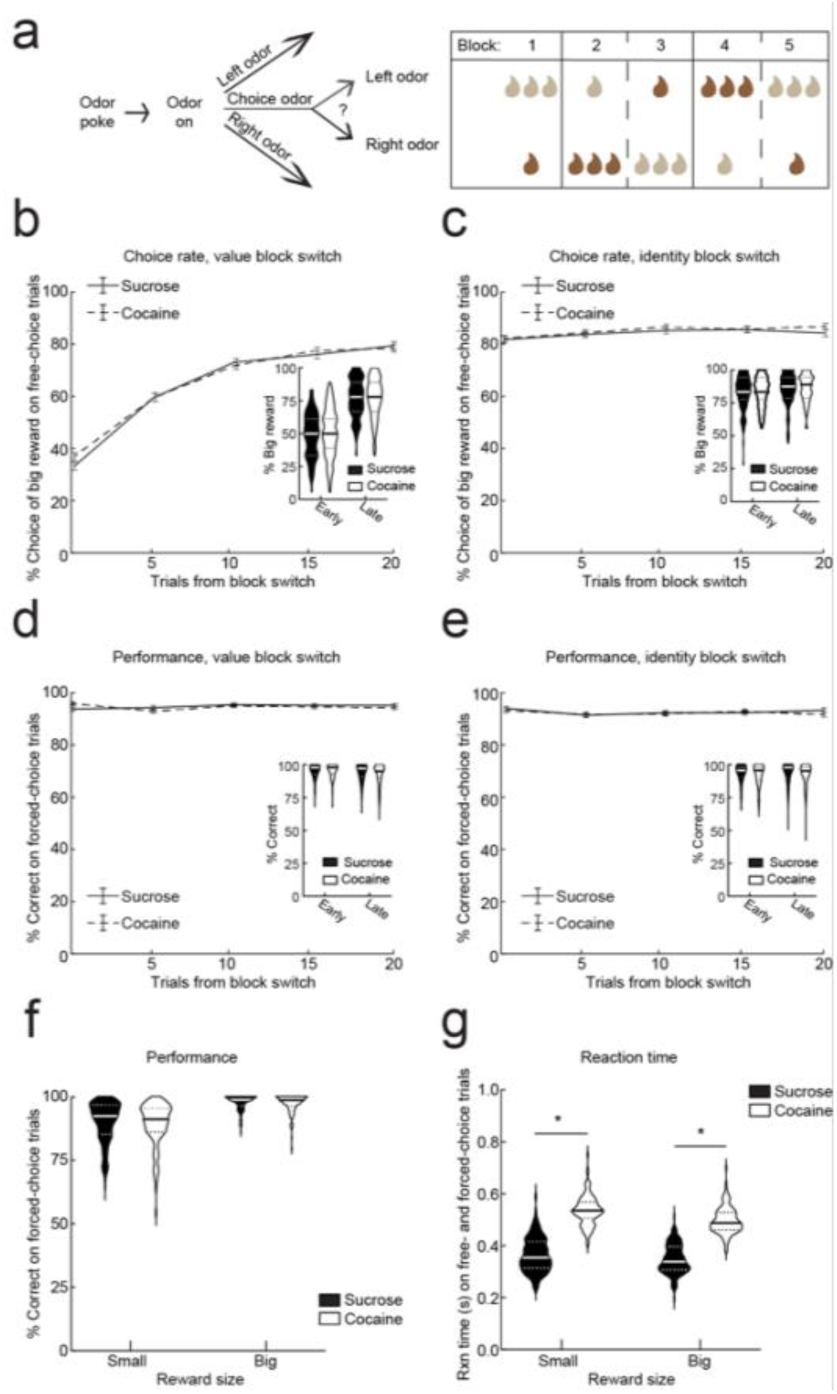
Task design and behavior. a) The decision-making task consisted of four unique blocks of trials defined by distinct response-reward contingencies. Instructional odors delivered to the central odor port indicated which action (go left, go right, go either direction) would be reinforced on each trial. On forced-choice trials, correct responses to the left or the right fluid well were reinforced with fluid delivery. On free-choice trials, responses to either fluid well resulted in fluid delivery. Fluid reinforcers delivered at each well differed in size and flavor and alternated following unsignaled block switches as shown in the example block sequence. b) The percent of big rewards chosen on free-choice trials was computed and aligned to value block switches (sucrose: solid line; cocaine: dashed line). Line indicates mean and error bars indicate SEM. *Inset*, violin plot show the percent of big rewards chosen on early versus late free-choice trials following value block switches (sucrose: solid black; cocaine: open). Solid lines in violin plots indicate medians and dots represent upper and lower quartiles; two-way ANOVA did not detect a significant treatment × time interaction: *P* = 0.5627, F(1,230) = 0.3360. c) The percent of big rewards chosen on free-choice trials was computed and aligned to identity block switches (sucrose: solid line; cocaine: dashed line). Line indicates mean and error bars indicate SEM. *Inset*, violin plot show the percent of big rewards chosen on early versus late free-choice trials following identity block switches (sucrose: solid black; cocaine: open). Solid lines in violin plots indicate medians and dots represent upper and lower quartiles; two-way ANOVA did not detect a significant treatment × time interaction: two-way ANOVA did not detect a significant treatment × time interaction: *P* = 0.9936, F(1,230) < 0.1. d) The percent of correct responses on forced-choice trials was computed and aligned to value block switches (sucrose: solid line; cocaine: dashed line). Line indicates mean and error bars indicate SEM. *Inset*, violin plot show the percent of correct responses on early versus late forced-choice trials following value block switches (sucrose: solid black; cocaine: open). Solid lines in violin plots indicate medians and dots represent upper and lower quartiles; two-way ANOVA did not detect a significant treatment × time interaction: *P* = 0.2433, F(1,230) = 1.368. e) The percent of correct responses on forced-choice trials was computed and aligned to identity block switches (sucrose: solid line; cocaine: dashed line). Line indicates mean and error bars indicate SEM. *Inset*, violin plot show the percent of correct responses on early versus late forced-choice trials following identity block switches (sucrose: solid black; cocaine: open). Solid lines in violin plots indicate medians and dots represent upper and lower quartiles; two-way ANOVA did not detect a significant treatment × time interaction: *P* = 0.4116, F(1,230) = 0.6768. f) The percent of correct responses on forced-choice trials for small (left) or big rewards (right) throughout the session (sucrose: solid black; cocaine: open). Solid lines in violin plots indicate medians and dots represent upper and lower quartiles; though both treatment groups demonstrated greater correct responses on big forced choice trials (significant main effect of size: *P* < 0.0001, F(1,230) = 292), a two-way ANOVA did not detect a significant treatment × size interaction (*P* = 0.6433, F(1,230) = 0.2151). g) Reaction times for small (left) and big rewards (right) on free- and forced-choice trials throughout the session (sucrose: solid black; cocaine: open). Solid lines in violin plots indicate medians and dots represent upper and lower quartiles; two-way ANOVA detected a significant treatment × size interaction: *P* < 0.0001, F(1,230) = 18.11, all asterisks indicate Bonferroni post-test results of *P* < 0.0001.

### Prior cocaine self-administration slowed decision making

During recording sessions, both sucrose- and cocaine-experienced rats preferred the big reward on free-choice trials, quickly adjusting their choice to the well producing the big reward over the first 20 trials following a value block switch (Fig. 1b), while exhibiting stable preference for the big reward following an identity block switch (Fig. 1c). Importantly, performance on forced-choice trials (i.e. selecting the correct well to get reward) remained high following both types of block switches (Fig. 1d, e). Additionally rats in both groups performed more accurately for the big reward on forced-choice trials and showed faster reaction times for the big reward on free- and forced-choice trials (Fig. 1f, g). Prior cocaine use did not impact any of these value-related measures, however cocaine-experienced rats did show significantly longer reaction times on free- and forced-choice trials for both big and small rewards as compared to sucrose-experienced rats (*P* < 0.0001 ; Fig. 1g).

To investigate whether prior cocaine experience altered neural activity in the OFC and PL, units were recorded in each of these regions (Fig. 2a). A previously established unsupervised hierarchical clustering method was used to divide unit populations in each region and treatment group into two clusters, one consisting of wide spikes (WS) and the other narrow spikes NS; ^45,46, 47^. OFC recordings in sucrose-experienced rats yielded a total of 1737 units of which 1545 were classified WS and 192 NS, and 1124 total units in cocaine-experienced rats with 970 WS and 154 NS (Fig. 2b). Session recordings in the PL of sucrose-experienced rats yielded 1080 units with 985 defined WS and 95 NS, and 1307 units in cocaine-experienced rats with 1209 defined WS and 98 NS (Fig. 2c). Proportions of classified WS and NS units per region were similar across treatment groups and were consistent with previous findings ^46–48^. Studies indicate that NS are likely putative interneurons, consisting of both fast-spiking and non-fast-spiking units, and WS are likely putative pyramidal units ^46, 47, 49–51^. In this study, we focus on WS unit populations.

**Figure 2.**
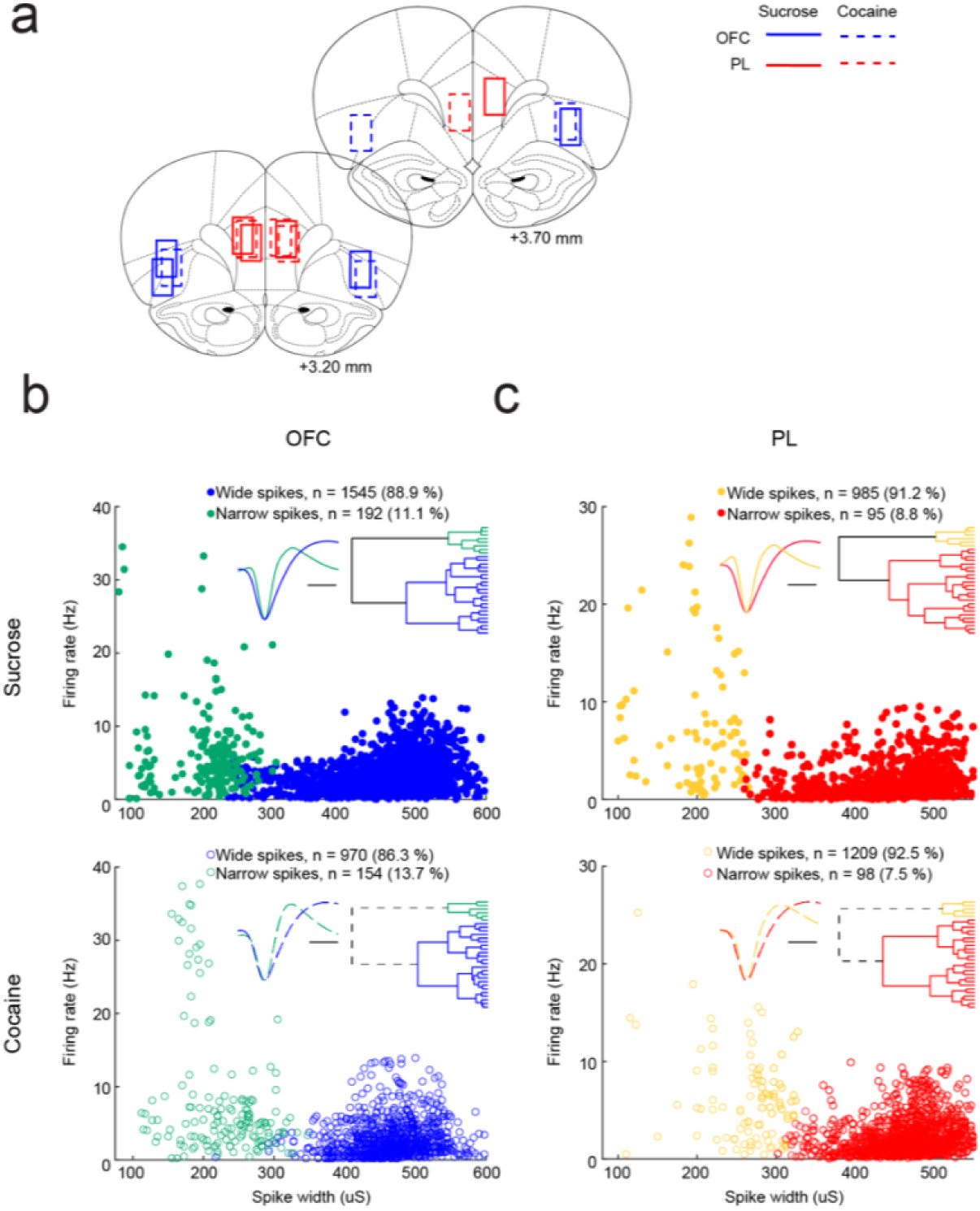
Histology and unit classification. a) Approximate location of neural recordings in the OFC and PL are indicated by boxes (sucrose: solid blue line; cocaine: perforated red line). b) Mean firing rates and spike widths for each OFC unit recorded from sucrose- (top; NS, 6.51 Hz, 205.70 μs; WS, 3.25 Hz, 461.17 μs) and cocaine-experienced rats (bottom; NS, 8.05 Hz, 229.69 μs; WS, 2.7 Hz, 465.08 μs) yielding two unique populations consistent with putative narrow-spiking interneurons (NS, green) and wide-spiking pyramidal cells (WS, blue). Dendrogram insets depict population clusters identified by the unsupervised cluster analysis. Wave insets show the average waveforms for classified unit populations. Scale bar; 100 µs. c) Mean firing rates and spike widths for each PL unit recorded from sucrose-(top; NS, 7.27 Hz, 207.50 μs; WS, 1.84 Hz, 446.22 μs) and cocaine-experienced rats (bottom; NS, 6.18 Hz, 265.1 μs; WS, 2.33 Hz, 468 μs) yielding two unique populations consistent with putative narrow-spiking interneurons (NS, yellow) and wide-spiking pyramidal cells (WS, red). Dendrogram insets depict population clusters identified by the unsupervised cluster analysis. Wave insets show the average waveforms for classified unit populations. Scale bar; 100 µs.

### Prior cocaine self-administration disrupted response dynamics of OFC and PL units

To examine whether prior cocaine experience altered the general response properties of OFC and PL WS units during performance of the decision-making task, free- and forced-choice trials were divided into five epochs related to each event comprising a trial (Supplemental Fig. 3), and units were sorted according to the time of maximum response across these epochs (Fig. 3a - d). This revealed a robust effect of prior cocaine use on firing rate dynamics in the OFC, with a significant decrease in the proportion of OFC WS units maximally active during the fixation and odor-sample epochs, and a significant increase in the proportion of units maximally active during the anticipation and consumption epochs in cocaine-versus sucrose experienced-rats (*P* < 0.0001; Fig. 3a, b). Prior cocaine use also caused changes in the firing rate dynamics in the PL, significantly increasing the proportion of maximally active PL WS units during the odor, movement and anticipation epochs, while decreasing the proportion maximally active during the consumption epoch (*P* < 0.0236; Fig. 3c, d).

**Figure 3.**
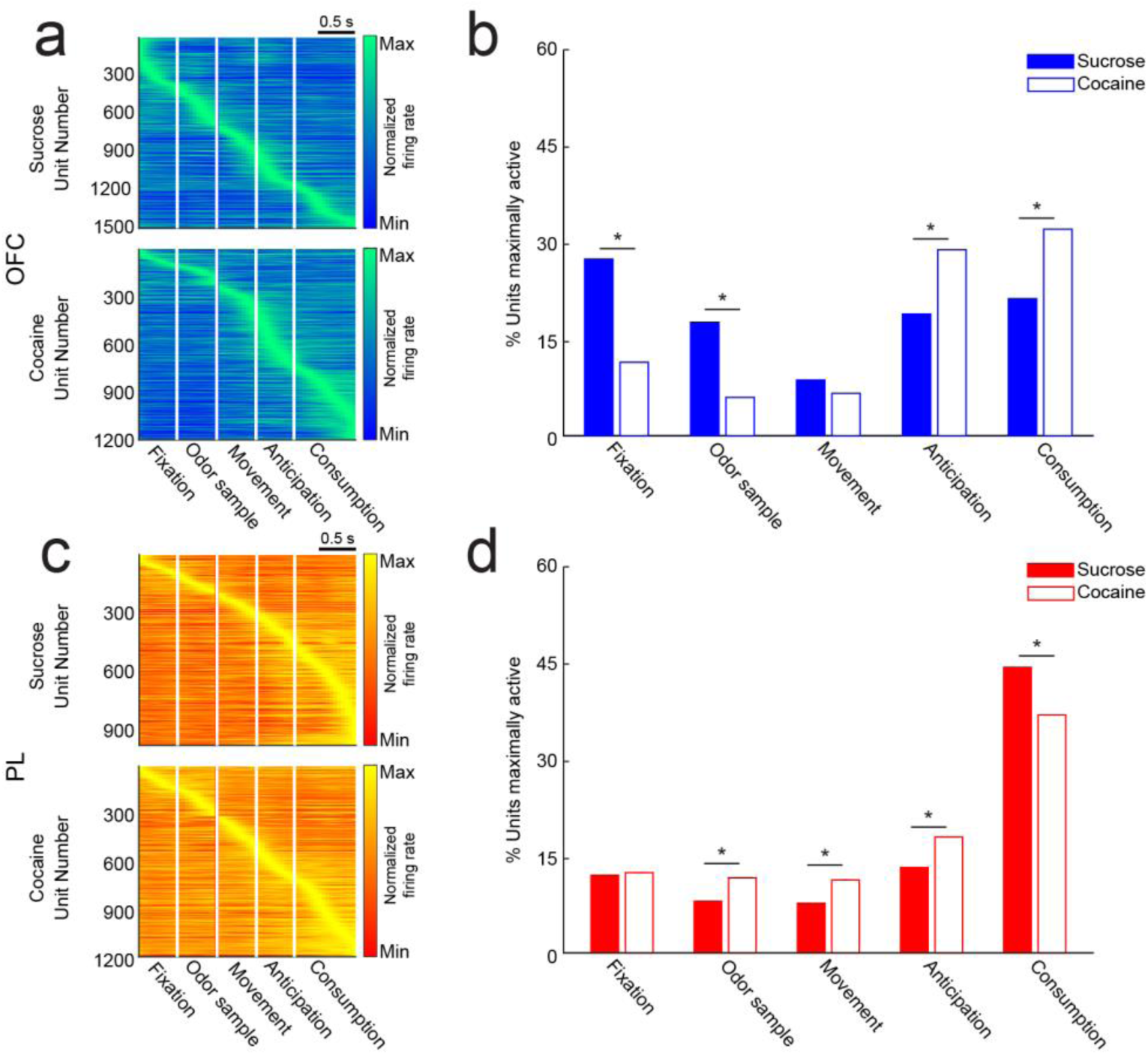
Time course of OFC and PL WS neural responses across treatment groups. a) Peri-event time histograms (PETHs) of peak-normalized firing rates in OFC WS recorded from sucrose-(top) and cocaine-experienced rats (bottom) aligned to each of the five task epochs. Each row shows the firing rate of a single unit using a color scale ranging from blue (zero) to green (peak) with time on the x-axis (each panel = 0.5 s or 0.8 s; bin size was 50 ms). b) Percent of OFC WS maximally active per epoch recorded from sucrose- and cocaine-experienced rats (sucrose: solid blue; cocaine: open blue). Z-test for population proportions, correct for multiple comparisons detected significant differences in the proportion of units maximally active across treatment groups during most epochs: fixation, *P* < 0.0001, z(2514) = 11.3921; odor sample, *P* < 0.0001, z(2514) = 7.0847; anticipation, *P* < 0.0001, z(2514) = −5.0833; consumption, *P* < 0.0001, z(2514) = −7.8152. All asterisks indicate z-test results of *P* < 0.0001. c) Peri-event time histograms (PETHs) of peak-normalized firing rates in PL WS recorded from sucrose- (top) and cocaine-experienced rats (bottom) aligned to each of the five task epochs. Each row shows the firing rate of a single unit using a color scale ranging from blue (zero) to green (peak) with time on the x-axis (each panel = 0.5 s or 0.8 s; bin size was 50 ms). d) Percent of PL WS maximally active per epoch recorded from sucrose- and cocaine-experienced rats (sucrose: solid red; cocaine: open). Z-test for population proportions, correct for multiple comparisons detected significant differences in the proportion of units maximally active across treatment groups during most epochs: odor sample, *P* = 0.0236, z(2193) = −2.8251; movement, *P* = 0.022, z(2193) = −2.854; anticipation, *P* = 0.0129, z(2193) = −3.0142; consumption, *P* = 0.0022, z(2193) = 3.5192. All asterisks indicate z-test results of *P* ≤ 0.0236.

### Prior cocaine self-administration reduced OFC selectivity for choice type

While the prior analysis demonstrated that cocaine experience disrupted the response properties of OFC and PL WS units, it did not address whether the encoded information was impacted. Similar to previous studies ^52–56^, we found that units from these cortical regions represented each of the four variables defining the trial and its outcomes – choice type (free or forced), response direction (left or right fluid delivery well), size (big or small), and flavor (chocolate or vanilla) (Fig. 4a, b).

**Figure 4.**
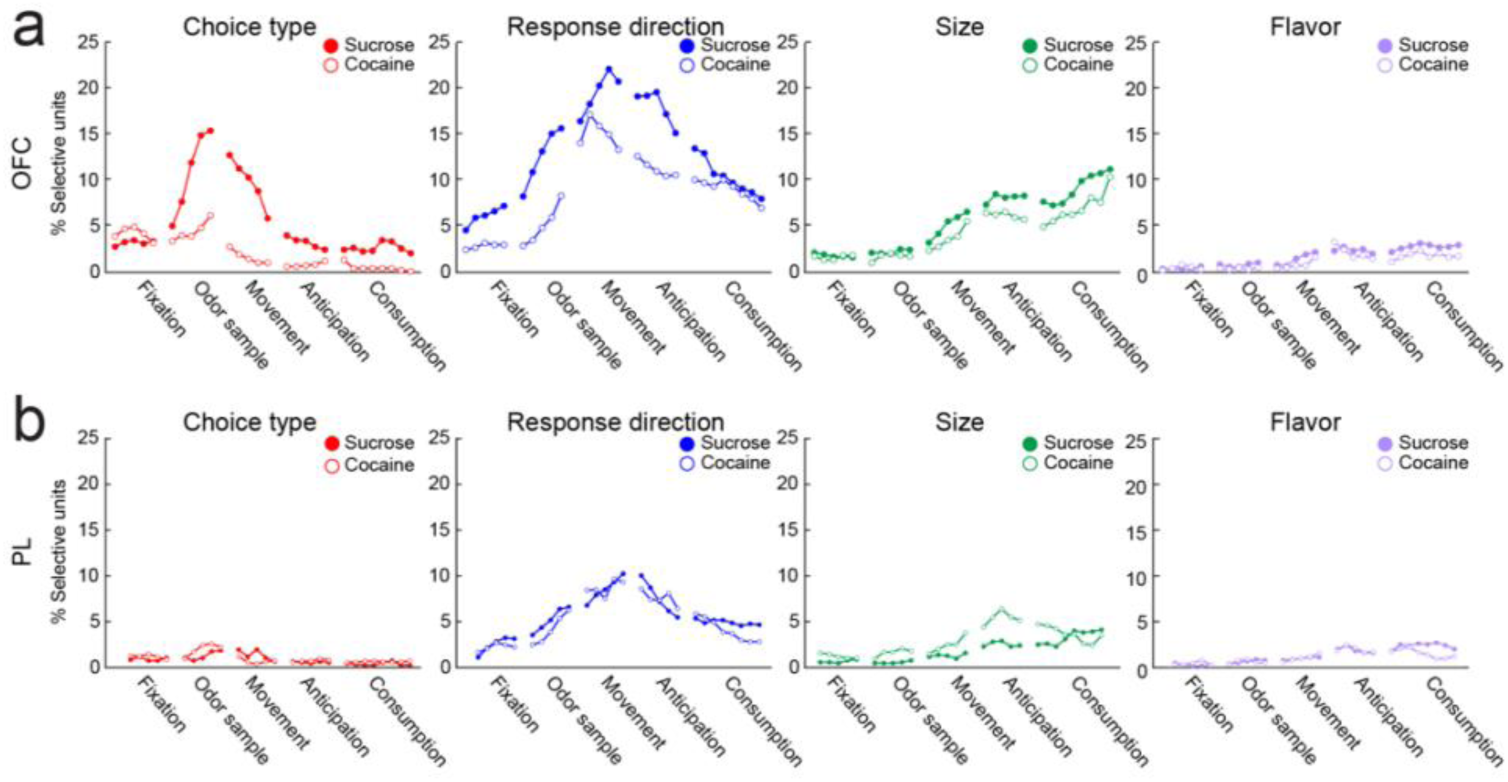
OFC and PL WS unit selectivity. a) Proportion of OFC WS units recorded from sucrose- and cocaine-experienced rats with activity significantly changed by trial-relevant dimensions of choice type (red), response direction (blue), size (green), or flavor (purple) throughout the trial (sucrose: solid; cocaine: open). The proportion of units with activity significantly modulated by these dimensions generally increased during or following the odor sample epoch (e.g., see ‘choice type’ panel), with the exception of the flavor dimension. The proportion of choice-type and response-direction selective units was lower in OFC WS activity recorded from cocaine-experienced rats compared to sucrose controls. Bin size used to fit models was 100 ms. b) Proportion of PL WS units recorded from sucrose- and cocaine-experienced rats with activity significantly changed by trial-relevant dimensions of choice type (red), response direction (blue), size (green), or flavor (purple) throughout the trial (sucrose: solid; cocaine: open). Though the proportion of units with activity significantly modulated by choice type was much lower in the PL (‘choice type’ panel) than the OFC (panel a, left), the proportion modulated by other trial-relevant variables generally increased during or following the odor sample epoch, with the exception of the flavor dimension. In contrast to OFC, the proportion of units selective for each dimension was generally not decreased in PL WS activity recorded from cocaine-experienced rats compared to sucrose controls, with in fact an increase in units with activity significantly modulated by size apparent during the anticipation epoch. Bin size used to fit models was 100 ms.

To compute selectivity for each of these single-dimension task variables across all WS units per cortical region recorded throughout performance of the task, we fit generalized linear models to the firing rate of each unit at every time bin throughout completed free- and forced-choice trials and plotted the percent of units selective for each individual variable (following Bonferroni-Holm correction for multiple comparisons) at each point (Fig. 4a, b). This showed that selectivity for most of the variables increased either during or following the odor sample epoch, the period in a trial that rats had knowledge about the choice type of the trial they were performing on, as well as the available actions and outcomes. While the proportion of OFC WS units selective for size and flavor was similar between treatment groups, the proportion selective for choice type and response direction was significantly lower in cocaine-experienced rats compared to sucrose-experienced rats (Fig. 4a). Interestingly, a distinct effect profile was observed in the PL, where the fraction of units selective for choice type, response direction and flavor were similar across treatment groups, and the fraction selective for size was significantly higher in the unit population recorded from cocaine-experienced rats as compared to sucrose controls (Fig. 4b).

### Prior cocaine self-administration decreased the strength of OFC choice type representations

To examine the strength with which task variables were encoded, we used d’ to quantify the separation of the distributions of correlation coefficients from within- (e.g., pairs of the same variables) and between-dimension conditions (e.g., pairs of opposing variables; Fig. 5a, b). That is, for each dimension of interest (e.g., choice type), we subtracted the average between-dimension distribution of correlation coefficients (e.g., each forced-choice-associated population vector correlated with each other free-choice-associated population vector) from the average within-dimension distribution of correlation coefficients (e.g., each free-choice-associated population vector correlated with each free-choice-associated population vector, or forced-choice with forced-choice) and then divided the values by the spread of the distribution (i.e., standard deviation) ^57^. For this, pseudoensembles were created by randomly choosing subsets of 100 units with replacement across all sessions. Pseudoensembles were used to create population vectors associated with each of the 16 unique trial variable combinations, which were then used to compute d’ for each of the four trial-related variables – choice type, response direction, size, and flavor. This was repeated 250 times for each region and treatment group. The analysis revealed that d’ values of choice type, response direction, size and flavor calculated using activity recorded from OFC WS in sucrose-experienced rats prior to odor delivery were low. Upon odor presentation, d’ of choice type increased (Fig. 5a, left), giving way to an increased d’ of response direction during the movement epoch (Fig. 5a, middle left), in turn giving way to an increased d’ of size during the anticipation and consumption epochs (Fig. 5a, middle right). Prior cocaine self-administration had differential effects on d’ values of the four dimensions with choice type significantly decreased (*P* = 0.0023; Fig. 5a, left), suggesting a weakening in the information being encoded by the OFC during trial performance.

**Figure 5.**
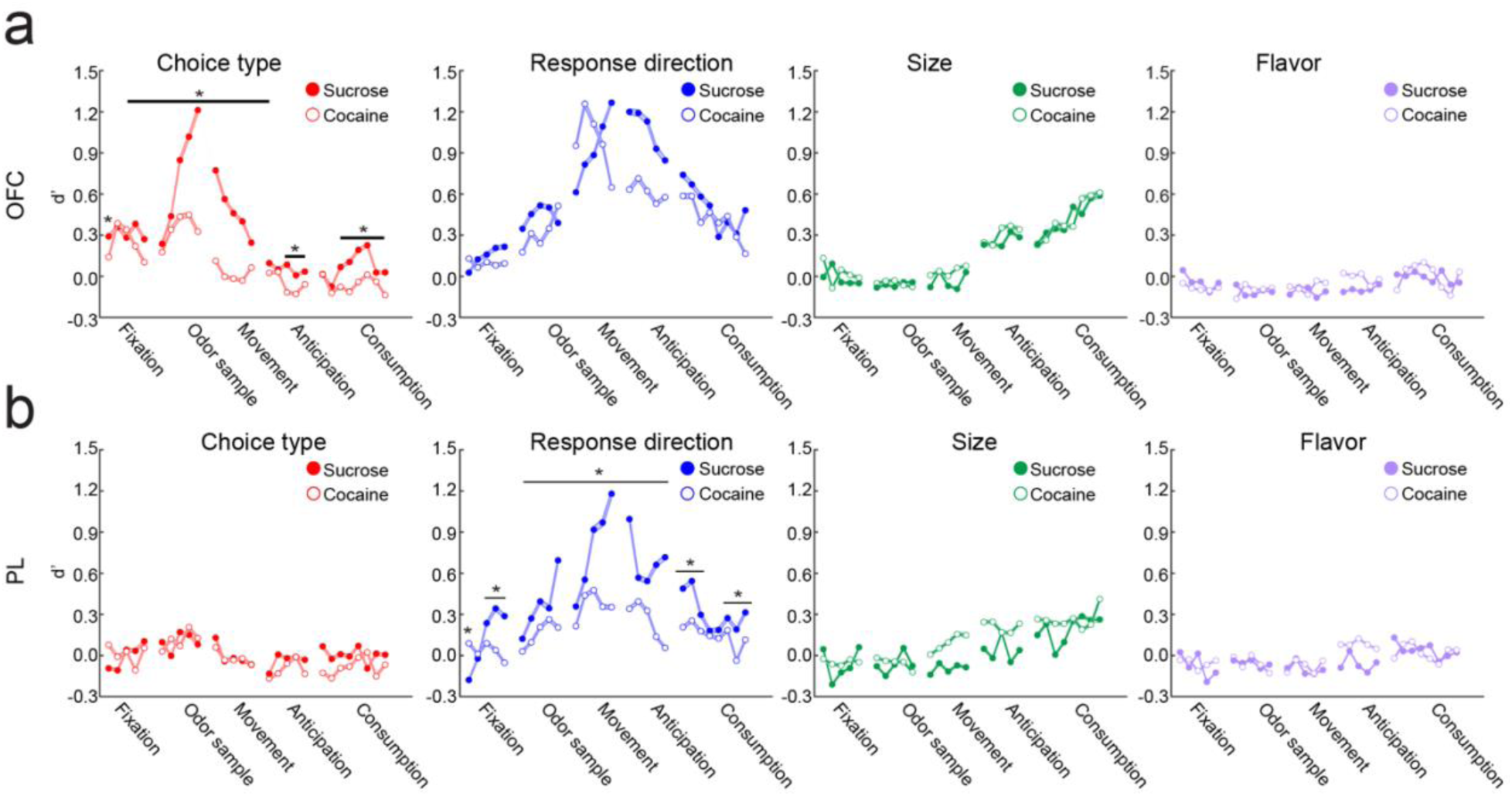
Strength of OFC and PL WS representations of trial dimensions over time. a) Mean d’ of trial-related variables using pseudoensembles of 100 OFC WS units randomly selected from pooled units recorded across all sessions from sucrose- and cocaine-experienced rats (sucrose: solid; cocaine: open). A two-way ANOVA detected a significant dimension × treatment interaction: *P* = 0.0146, F(3,216) = 3.584; Bonferroni post-tests showed a significant decrease in d’ of choice type (*P* = 0.0023). A follow-up two-way ANOVA on d’ of choice type detected a significant time × treatment interaction: *P* < 0.0001, F(27,13944) = 162.9, all asterisks indicate Bonferroni post-test results of *P* ≤ 0.0359. Bin size was 100 ms; shading indicates the SEM in of d’. b) Mean d’ of trial-related variables using pseudoensembles of 100 PL WS units randomly selected from pooled units recorded across all sessions from sucrose- and cocaine-experienced rats (sucrose: solid; cocaine: open). A two-way ANOVA detected a significant dimension × treatment interaction: *P* < 0.0001, F(3,216) = 13.58; Bonferroni post-tests showed a significant decrease in d’ of response direction (*P* < 0.0001). A follow-up two-way ANOVA on d’ of choice type detected a significant time × treatment interaction: *P* < 0.0001, F(27,13944) = 396.4, all asterisks indicate Bonferroni post-test results of *P* ≤ 0.0042. Bin size was 100 ms; shading indicates the SEM of d’.

The same analysis performed using pseudoensembles of PL WS units recorded from sucrose-experienced rats revealed that the highest d’ value was response direction, while those of choice type, size, and flavor were low through most epochs of the trial (Fig. 5b). During the trial, d’ value of response direction became particularly high at the end of the odor sample epoch and through the movement epoch. Prior cocaine self-administration had differential effects on d’ values of the four dimensions with response direction significantly decreased (*P* < 0.0001; Fig. 5b, second from left).

### Prior cocaine self-administration altered the organization of OFC multidimensional hierarchical representations

We next explored how task-relevant information was integrated and structured at the population level in the OFC and PL. We used a representational similarity analysis (RSA) ^57, 58^ to look at the geometry of the neural activity space across the multiple dimensions characterizing task trials. To compute the similarity of population level ensemble representations of different trial dimensions for each cortical region, we calculated the median z-normalized firing rate for each WS unit for all 16 unique choice type x response direction x size x flavor trial combinations during the odor sample epoch and constructed a population vector for each combination based on normalized rates. The relationships between ensemble representations were then measured by computing the correlation coefficient between population vectors for every pair of trial-related variable combinations. To visualize the organization, or hierarchy, of information represented by WS unit populations per cortical region during the odor sample epoch, dendrograms were constructed by grouping correlation values into clusters (Fig. 6a, b).

**Figure 6.**
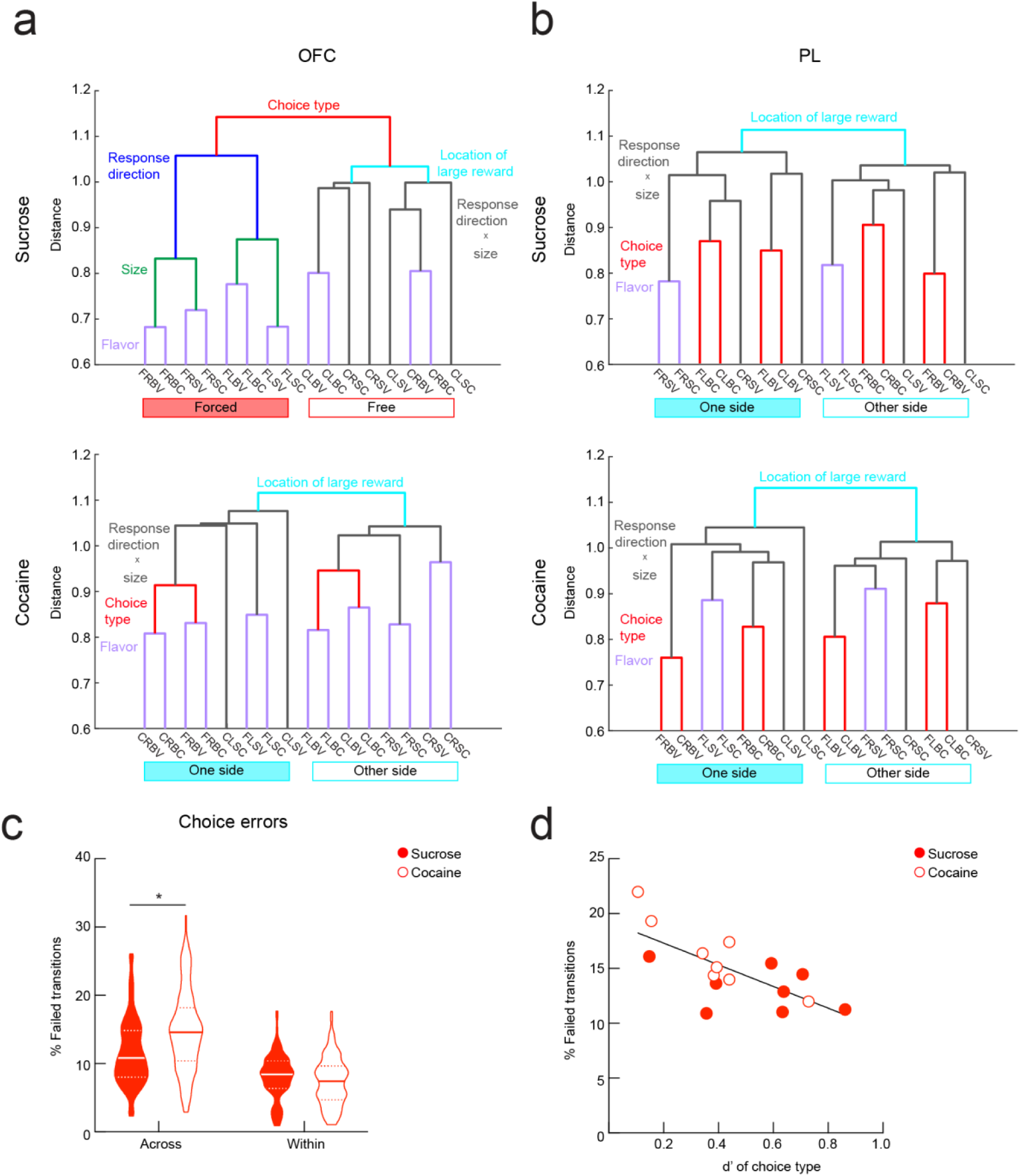
OFC and PL WS population ensemble hierarchical representations during odor sampling. a) Dendrograms from OFC WS population ensembles recorded from sucrose- and cocaine- experienced rats (top and bottom, respectively), showing the format of information represented during the odor sample epoch as a function of correlation of population vectors along various trial-related dimensions. Visualization of dendrograms shows that prior cocaine experience alters the format of information represented in the OFC. Note that in the top panel, the most separable dimension is choice type, whereas in the bottom panel the most separable dimension is the location of the large reward. See main text for further description. b) Dendrograms from PL WS population ensembles recorded from sucrose- and cocaine-experienced rats (top and bottom, respectively), showing the format of information represented during the odor sample epoch as a function of correlation of population vectors along various trial-related dimensions. Visualization of dendrograms shows that prior cocaine experience does not change the formation of information represented in the PL. Note that in both top and bottom panels the most separable dimension is location of the large reward. See main text for further detail. c) Across-choice transition errors on the left of the graph are defined as either (1) an incorrect response on a forced-choice trial following a free choice or (2) choice for the small reward on a free-choice trial following a forced choice. Within-choice transition errors on the right of the graph are defined as either (1) an incorrect response on a forced-choice trial following a forced choice or (2) choice for the small reward on a free-choice trial following a free choice. Sucrose: solid red; cocaine: open red. Two-way repeated-measures ANOVA revealed a significant behavioral error × group interaction: *P* < 0.0001, F(1, 230) = 27.31. Post-tests showed that cocaine-experienced rats made a greater fraction of across-choice transition errors: *P* < 0.0001, t(460) = 5.192, whereas both groups rats made a similar fraction of within-choice transition errors: *P* = 0.42, t(460) = 1.255. The asterisk indicates a Bonferroni post-test result of *P* < 0.0001. Solid lines indicate medians and dots represent upper and lower quartiles. d) Scatter plot of OFC WS d’ of choice type values computed using pseudoensembles from sucrose- and cocaine-experienced rats (sucrose: solid red; cocaine: open red) during the odor sample epoch in bins of two recording sessions across the experiment against choice transition errors averaged over corresponding sessions. Generalized linear models fit to across-choice-type transition errors to examine how well behavior could be explained by d’ of choice type showed that d’ significantly predicted task performance: *P* = 0.0085, t(13) = −2.6292. Line indicates the best fit of pooled sucrose and cocaine session data.

The dendrogram produced using the OFC WS unit population recorded from sucrose-experienced rats revealed that the most separable representation was choice type (free- or forced-choice), under which the hierarchical structure of information differed. On forced-choice trials, when rats were required to respond to the fluid delivery well signaled by the odor cue, response direction was the most separable representation, followed by size and then flavor. However, on free-choice trials, when rats chose their preferred fluid delivery well, trial types first grouped based on the location of the large reward, followed by combinations of response direction and size, and then flavor (Fig. 6a, top). Prior cocaine experience altered the format of information encoded by the OFC WS unit population (Fig. 6a, bottom).

A much differently formatted structure was generated using the PL WS unit population recorded from sucrose-experienced rats, with the greatest separable representation being the location of the large reward (i.e., trials types were clustered based on whether they occurred before or after the value switch). Following this, a more ambiguous structure remained, with the next most separable features being combinations of response direction, size, and flavor. In stark contrast to the organization of OFC, choice type was relegated to a minor role (Fig. 6b, top).

Dendrograms generated using activity recorded from cocaine-experienced rats revealed a very similar organization in PL cortex, suggesting that prior cocaine experience does not alter the format of information represented by the PL WS unit population (Fig. 6b, bottom), whereas the organization in the OFC changed dramatically. Rather than being dominated by a division between free- and forced-choice trials, the organization in the OFC of cocaine-experienced rats became much more similar to that observed in PL, becoming dominated by the location of the large reward, followed by an ambiguous division of response direction and size, choice type, and flavor (Fig. 6a, bottom).

### Representations of choice type in OFC unit population correlates with task performance

Because prior cocaine experience decreased the prevalence of choice-type selective units (Fig. 4 a), the strength of choice type representations (Fig. 5a), and the structure of information in the OFC reflecting the difference between free- and forced-choice trials (Fig. 6a), we next questioned whether differences in representational strength of choice type might be related to differences in behavior across treatment groups. We postulated that if the normal separation of choice type in the neural activity space in OFC was behaviorally relevant then there might be behavioral differences when transitions were required between forced- and free-choice trials (i.e., choice type transition), wherein that weaker representations would correspond to less optimal behavior.

To test this, we first examined the ability of rats to make optimal responses following different choice-type transitions. Across-choice transition errors were defined as either (1) an incorrect response on a forced-choice trial following a free choice trial or (2) choice for the small reward on a free-choice trial following a forced choice. Within-choice transition errors were defined as either (1) an incorrect response on a forced-choice trial following a forced choice or (2) choice for the small reward of a free-choice trial following a free choice. Prior cocaine use was associated with greater across-choice but not within-choice transition errors (*P* < 0.0001; Fig. 6c). To test whether this behavioral finding was related to the loss of separation between free- and forced-choice trials in the neural activity space in OFC, we next fit a generalized linear model to across-choice transition errors to look at how well behavior could be explained by the d’ of choice type representations during the odor sample epoch, focusing on this dimension because it was the most affected by cocaine in the OFC (Figs. 4a, 5a, and 6a). The results showed d’ of choice type was significantly correlated with the rate of across-choice transition errors (*P* = 0.0085254; Fig. 6b).

### Prior cocaine self-administration decreased encoding of latent information by OFC

Finally, we examined whether the geometry of neural representations in OFC and PL was affected by previous cocaine experience. We computed the shattering dimensionality of OFC and PL ensembles—a measure of how separable arbitrary groupings of representations are from one another. Higher shattering dimensionality implies a high-dimensional neural representation from which downstream structures could flexibly read out a wide range of information, including information that was orthogonal or even contradictory to task-relevant information at the time of learning.

For this analysis, we took firing rates during the odor sample epoch on the task’s eight forced-choice trial types (which were defined by unique response direction × size × flavor mappings), grouped activity on all balanced trial type combinations possible (i.e., 35 unique ways of evenly pairing eight trial types – the three corresponding with known task-related variables, such as response direction, size, and flavor were excluded, with the other 32 arbitrary pairings remaining as latent variables), and used cross-validated support vector machines (SVM) to classify trial type from neural activity on each of the 32 trial-type pairings ^59;^ Fig. 7a, c. Free-choice trials, in which rats strongly preferred big rewards, were excluded to ensure similar numbers of trials in all groupings.

**Figure 7.**
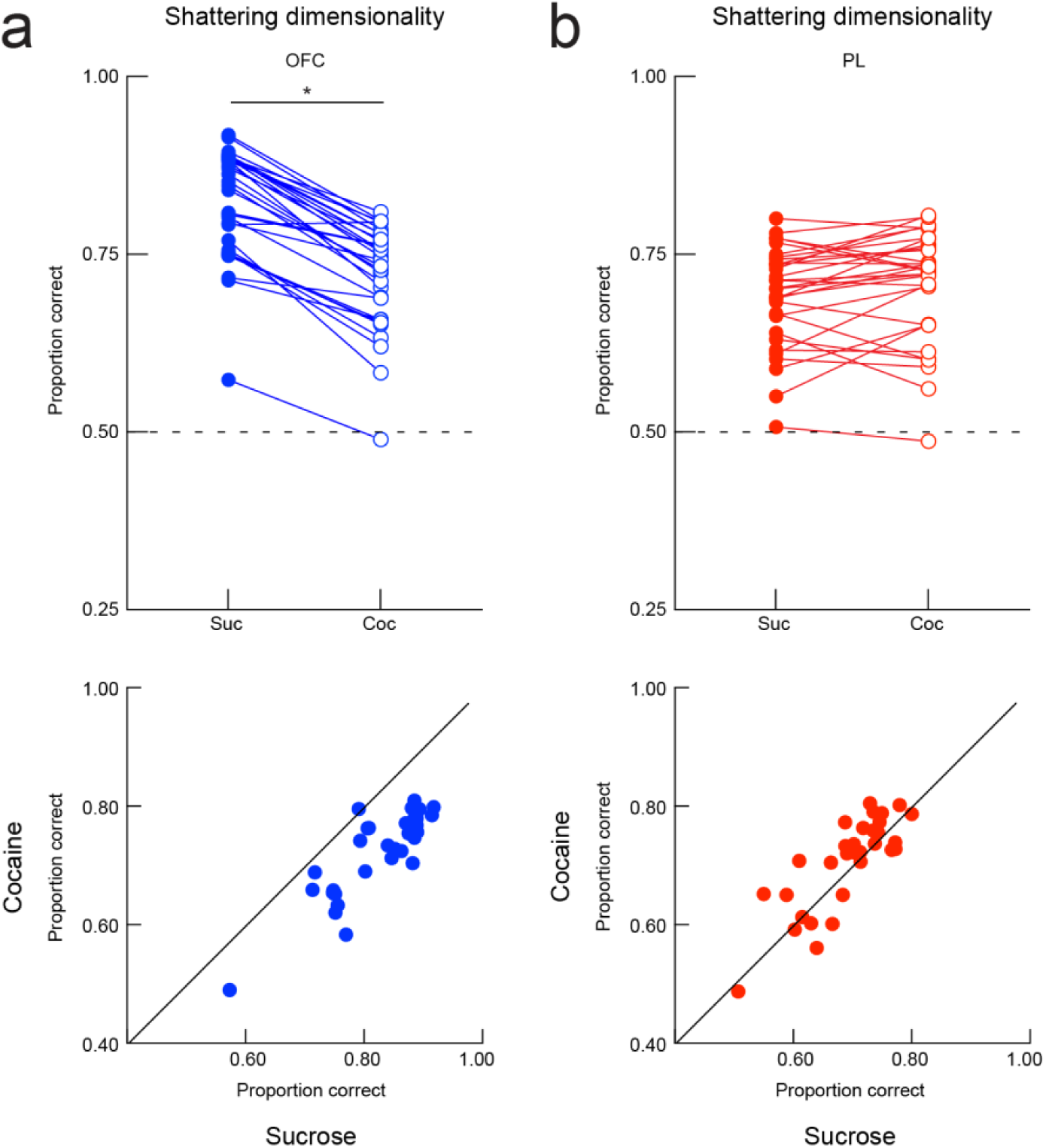
Representation of latent information by OFC and PL WS during odor sampling. a) Top, proportion of correct classification for the 32 dichotomies corresponding to latent information variables for OFC WS pseudoensembles drawn from sucrose- and cocaine-experienced rats (sucrose: solid blue; cocaine: open blue) during odor sampling. Paired Student’s t-test detected significantly decreased values in the cocaine group: *P* < 0.0001, t(30) = 13.76. The asterisk indicates a result of *P* < 0.0001. Bottom, proportion of correct classification for the 32 dichotomies corresponding to latent information variables for OFC WS pseudoensembles from sucrose-experienced rats plotted against those from cocaine-experienced rats. Visualization of the data showed that more values were biased toward higher sucrose classification and fell below the perfectly correlated line b) Top, proportion of correct classification for the 32 dichotomies corresponding to latent information variables for PL WS pseudoensembles drawn from sucrose- and cocaine-experienced rats (sucrose: solid blue; cocaine: open blue) during odor sampling. Paired Student’s t-test did not detect different values across treatment groups: *P* = 0.0928, t(30) = 1.736. Bottom, proportion of correct classification for the 32 dichotomies corresponding to latent information variables for PL WS pseudoensembles drawn from sucrose-experienced rats plotted against cocaine-experienced rats. Visualization of the data showed that many values fell along the perfectly correlated line.

Classification accuracy exceeded chance performance in both OFC and PL (Fig. 7a, b). Shattering dimensionatly was greater in OFC than PL (Fig. 7a, b). Further, cocaine experience reduced shattering dimensionality in OFC, with nearly every single variable grouping showing lower classification accuracy in the cocaine group (*P* < 0.0001; Fig. 7a). In contrast, cocaine experience did not affect the shattering dimensionality of PL population representations (Fig. 7b). These results show that cocaine experience profoundly alters the geometry of ensemble representations in OFC, but not in PL.

## Discussion

Here, we found that prior cocaine experience significantly altered the strength and structure of task-relevant and latent information in the OFC and that these changes were related to less optimal decision making. Single-unit-level regression analyses looking at the encoding of four known task-related variables – choice type, response direction, size, and flavor – showed that prior cocaine experience significantly decreased the number of choice-type and response-direction selective units in OFC. At the population level, representational similarity analyses looking at the encoding of multiple dimensions of the same known task-related variables revealed that prior cocaine experience changed the structure of information represented in the OFC, making it more focused on size or value and eliminating distinction in the organization of information between free and forced-choice trials. This change in the importance of choice type was also reflected in d’ analyses, which found a significant weakening of choice-type representations in OFC after cocaine use. These changes were related to worse task performance at transitions between free- and forced-choice trials. Finally, examination of the encoding of latent information, using shattering dimensionality, showed that prior cocaine use also decreased the representation of latent information in the OFC. By contrast, cocaine had comparatively marginal effects on the PL, with prior drug use increasing the prevalence of size-selective units and weakening the strength of response direction representations during the task.

### Prior cocaine self-administration preferentially impacts representations in OFC

The effects of cocaine were more dramatic in the OFC than in PL. This difference may be related to the task requirements. Though both cortical regions are important for flexible decision-making, previous work highlights the specific importance of the OFC for adaptive behavior when reward value or contingencies change, with lesions resulting in the inability to quickly adjust behavioral responding following changes in reward contingencies ^17, 60^, as in reversal ^43, 61, 62^, or after reinforcer devaluation ^17, 18^. By contrast, the PL is required during changes in the attentional demands of a task, for rats to express an instrumental response reflective of the current optimal goal value, and for using environmental stimuli to resolve response conflict^25, 63, 64^, as is the case when switching between response and strategy sets ^65–68^. Our task places little demand on changes in strategy or attentional set, and the conditions generally have similar value and thus similar attentional demands, across choice types and response directions. Thus, the differences in the representations of task structure between the two areas and the generally reduced activity in PL – and modest effect of cocaine in PL – may reflect this asymmetry in the emphasis of the task on OFC versus PL dependent processes.

Alternatively, however it is possible that drugs – at least cocaine – do have an outsized effect on processing in OFC. This would be consistent with repeated findings that use of addictive drugs impair OFC-associated functions across species^69, 70^. This includes behavior and learning in settings such as reversal learning^4, 6, 7, 9, 10, 38^, devaluation^11, 12^, Pavlovian over-expectation^71^, and sensory preconditioning^15, 72^, in which the OFC is proposed to mediate the use of cognitive maps to support inference^73^. Conversely limited effects on medial prefrontal processing would be consistent with recent work showing that comparable instrumental tasks, known to depend on medial prefrontal regions, are much less sensitive to drug effects^74^. Thus, OFC-dependent cognitive mapping and inference may be selectively impacted by drug use, an effect that would have significant implications for the control of behavior and learning.

### Prior cocaine self-administration reorganizes and reduces the complexity of task relevant representations in OFC

Cocaine did not eliminate selective activity in OFC or even substantially change the numbers of neurons active in the task or in particular epochs for the most part; instead there was a reorganization of the way that the task relevant variables – choice type, response direction, size, and flavor – were represented. While the neural activity space in controls exhibited a major separation between free and forced choice trials, the activity space in OFC in cocaine-experienced rats was primarily organized by value, with choice type becoming a minor variable and size becoming the main organizing principle, followed by response direction. In addition, while the organization of the other variables under the top line variable of choice type was quite different in controls, the organization of the more minor variables under the top line variable of size was much more ordered in the cocaine-experienced rats. One way to view this is that the activity space in controls represented free and forced choice trials as if they were different tasks, with the associated complexity, whereas after cocaine use, the activity space became much simpler and based simply on locating the large reward. This result is consistent with modeling work, suggesting that normal rats learn this task in a way that suggests separate representations for the two trial types^75^. Maintaining different task representations can be beneficial, particularly when it is necessary to respond differently; consistent with this, the neural separation based on choice type was inversely correlated with the propensity to make errors – select the less-valuable well – when a free choice trial followed a forced choice trial or vice versa.

### Prior cocaine self-administration impoverishes representation of latent information in OFC

We assessed the impact of prior cocaine use on the representation of latent information in the OFC formally by using the method of calculating shattering dimensionality, developed and highlighted in recent work on prefrontal cortex and hippocampus as an unbiased way to assess representation of latent information ^59^. This approach quantifies the ability to decode nonsense groupings of trial sets in the neural activity space. In our design, across 32 such pairs, the average decoding by ensembles of OFC neurons in controls was nearly 80%, an astonishing level of accuracy given that the OFC is often proposed to represent only meaningful or value-relevant information and these groupings were at best orthogonal to such information in the task. As noted in the original work ^59^, the ability to represent task relevant information while not losing sight of other features allows a network to remain sensitive to information that may become important at another time or place. This ability may be a core function of prefrontal cortex, and within prefrontal cortex, the OFC may be particularly specialized for recognizing, tracking, and even preserving such latent states ^44, 76–79^. Indeed one way to think of devaluation, rapid reversal learning, and many other OFC-dependent tasks is that they require the use of information concerning aspects of predicted events that was irrelevant or of only potential significance at the time of learning. According to this interpretation, the critical contribution of the OFC is not representing value or even calculating it on the fly but rather is in keeping information of uncertain relevance discriminable within the state space ^80, 81^.

In the current study, prior cocaine use caused a marked loss of the ability to represent such information, at least as assessed by shattering dimensionality; OFC neurons recorded from cocaine-experienced rats performed basically at chance at decoding the nonsense trial sets, as if they had completely collapsed these hidden variables. Such simplification is consistent with the other findings, particularly the reorganization of the activity space based on value and the compression of free- and forced-choice trial types, both places where interesting but arguably irrelevant or low-relevance information has been jettisoned.

The loss of the ability to maintain latent information of normal complexity within higher cortical decision-making areas as a result of drug use would have enormous implications, essentially favoring pursuit of previously experienced rewards in context, particularly ones that are reliable and immediate, over the ability to integrate experience gained out of context or with events that are associated with delayed, improbable, rare, or even anecdotal significance. In some regards, this is an apt description of substance use disorder. Individuals suffering from SUD pursue drug rewards, which tend to be reliable and immediate, particularly in context, while showing less sensitivity to non-drug outcomes – both alternative rewards and possible punishments – which are typically rare, often removed from actual drug use, or learned about anecdotally or in other contexts.

Future studies examining the individual differences that may contribute to SUDs are necessary. Because the current study utilizes population vector analyses, made possible by combining all units recorded across all rats and all sessions per region and treatment group, it did not allow us to study between-rat differences in neural encoding and how it relates to behavior. To this end, recent work using reinforcement learning models to examine drug-naïve rat behavior on a similar task suggests that there are individual differences in representational learning ^75^, warranting a close examination of individual differences on the task and, further, whether cocaine intake impacts it. More subjects in combination with the ability to record higher numbers of simultaneous units in the future may allow for these detailed investigations, perhaps providing finer resolution to the relationship of representations and behavior in control animals and how this relationship is altered by prior cocaine experience.

Another avenue warranting further investigation is whether OFC manipulations (e.g., via optogenetics or pharmacogenetics) can rescue the behavioral deficits observed in our paradigm. Our lab has shown that local optogenetic stimulation of the OFC is sufficient to correct cocaine-associated behavioral impairments in one OFC-dependent task ^71^. Using a similar approach within the context of the present study would further strengthen the role of the OFC in representing both observable and hidden states. Lastly, as discussed above, to get a better understanding of the effect of prior cocaine experience on representations in the OFC and PL, designing a task that incorporates contingency reversals as well as intra- and extra-dimensional strategy shifts will be necessary. Continuing to characterize the fundamental, nuanced roles of these brain regions in behavior and how they are affected by drugs of abuse will help to better understand the neuropathological underpinnings of addiction and develop more effective targeted treatment strategies in the future.

## Author Contributions

LEM, AMW, and GS designed the experiment, and LEM, MJS, and CK conducted the experiment (surgery, behavioral training, unit recording, and post-mortem tissue processing). LEM analyzed the resultant data, with input from AMW and GS, and LEM, AMW, and GS interpreted the data and wrote the manuscript.

## Acknowledgments

This work was supported by the Intramural Research Program at the National Institute on Drug Abuse. The opinions expressed in this article are the authors’ own and do not reflect the view of the NIH/DHHS. The authors have no conflicts of interest to report.

## Data & Code Availability

The dataset and all scripts used in this study will be made available upon publication.

## Online Methods

### Subjects

Eight male Long-Evans rats (250-300 g, Charles River Labs), aged approximately three months, were subjects for this experiment. During the odor-guided choice task training and recording sessions, rats received water *ad libitum* for 10 minutes per day and *ad libitum* food. During self-administration, rats were food deprived to 85% of initial weight with *ad libitum* water. Self-administration and odor task testing were performed during the light phase. All experimental procedures complied with Institutional Animal Care and Use Committee of the US National Institute of Health guidelines.

### Surgical methods for jugular catheter and electrode implantation

All surgeries were performed under aseptic conditions as previously described ^82^. Rats were randomly assigned to cocaine or sucrose self-administration groups. Rats used for cocaine self-administration (n = 4) received chronic indwelling jugular catheter implants; rats used for sucrose self-administration (n = 4) received sham surgeries in which the jugular vein was exposed but no catheter was implanted. Rats recovered for seven days before self-administration began. During recovery and self-administration, catheters were flushed daily with a cocktail of enrofloxacin and heparinized saline to maintain patency. Following self-administration and training on the decision-making task, recording electrodes composed of drivable bundles of sixteen 25-µm diameter NiCr wires electroplated to an impedance of ∼200 kΩ were implanted into the PL of one hemisphere and the OFC of the opposite hemisphere in each rat (PL: 2.9 mm AP, ±0.7 mm ML, 3.5 mm V ^83^; OFC: 3.0 mm AP, ±3.2 mm ML, 4.0 mm V ^56^; relative to bregma and dura). The hemispheres of the implanted regions were counterbalanced across rats. Rats were allowed two weeks to recover and then returned to the decision-making task for one week of reminder training and subsequent recording sessions. Electrodes were advanced following each recording session to capture new units.

### Self-administration

The self-administration procedure was similar to that described previously ^71^. Briefly, rats were trained to self-administer intravenous cocaine-HCl (0.75 mg/kg/infusion; n = 4) or oral sucrose (10%, wt/vol; n = 4) under a fixed ratio 1 schedule in 3 h sessions over 14 consecutive days.

### Odor-guided choice task

The decision-making task was similar to that described previously ^44^. Task training and recording was performed using chambers equipped with two fluid wells and an odor port. Self-paced sessions began with the illumination of a house light, with trials initiated by a nose poke in the odor port. Upon odor port entry, rats held for a 500-ms fixation period, which was followed by a 500-ms presentation of one of three instructional odors that remained constant throughout the experiment. Following odor presentation, rats withdrew from the odor port and indicated their choice by entering either the left or right fluid well within 3000 ms. Upon fluid well entry, rats held for 500 ms before fluid reward delivery began. Once rats consumed the reward and withdrew from the fluid well, the house light was extinguished, and the trial ended. If rats withdrew prematurely at any point prior to fluid delivery, the trial was aborted, and the house light turned off. Two instructional odors specified a forced-choice reward delivery at either the left or the right fluid well. The third odor indicated a free choice trial. Presentation of odors occurred in a pseudorandom sequence with forced-choice right/left odors delivered equally on 65% of trials and free-choice odors delivered on 35% of trials.

Reward outcomes differed in flavor identity (vanilla or chocolate milk) and size (small [single 0.05 mL drop] or big [three 0.05 mL drops]), with opposite flavor and size outcomes being delivered at opposing fluid wells. Sessions consisted of a short initial block and followed by four blocks of approximately 60 trials each. Response-reward contingencies were consistent for blocks of trials but switched across blocks; block switches were not explicitly signaled and their timing varied randomly to prevent anticipation. Rats were trained before electrode implantation until they performed forced-choice trials with greater than 70% accuracy and completed all blocks. Following electrode implantation and recovery, rats received an additional week of reminder training to increase task performance and acclimation to recording cables.

### Single-unit recording

Neural activity was recorded using Plexon Multichannel Acquisition Processor Systems (Plexon Inc.) interfaced with a training chamber. Electrode signals were amplified 20x via operational head stages on electrode arrays and then passed to differential preamplifiers where they underwent an additional 50x amplification and filtration at 150-9000 Hz ^44, 82^. From there, signals were passed to multichannel acquisition processors where they underwent an additional 250-8000 Hz filtration, 40 kHz digitization, and final 1-32 x amplification. Signal waves were derived from active channels with event timestamps documented by the behavioral program.

### Experimental Design and Statistical Analyses

#### Behavioral epochs

Task trials were divided into five epochs either 0.5 s or 0.8 s in length (Supplemental Fig. 3). The fixation epoch began at time of odor port entry, which required rats to hold their snouts in the odor port, and ended after 0.5 s, immediately prior to odor cue delivery. The odor sample epoch began at the time of odor onset and ended after 0.5 s when odor cue delivery ended. Once the odor cue presentation ended, rats chose between the two fluid delivery wells. The movement epoch comprised 0.5 s immediately prior to entry into the fluid delivery well. The anticipation epoch began at the time of fluid delivery well entry and ended after 0.5 s, immediately prior to outcome delivery. The consumption epoch began at the time of outcome delivery and ended after 0.8 s.

#### Spike sorting and unit type classification

Neural activity was stored and manually sorted into putative single units using Offline Sorter (Plexon Inc.). Files were processed using NeuroExplorer (Nex Technologies) to extract timestamps and further analyzed using MATLAB (The MathWorks, Inc.). Orbitofrontal and prelimbic cortical populations were classified as wide waveform units (wide spike; WS) and narrow waveform units (narrow spike; NS) using a hierarchical unsupervised cluster analysis as previously described ^45, 46, 83^. Scatter plots of spike waveform widths (µs) and firing rates (Hz) show resulting clusters per region and treatment group (OFC: Fig. 2b; PL: Fig. 2c). Of the 1737 units recorded from the OFC of sucrose-experienced rats, 1545 were classified as WS and 192 were classified as NS. Of the 1124 units recorded from the OFC of cocaine-experienced rats, 970 were classified as WS and 154 were classified as NS. Of the 1080 units recorded from the PL of sucrose-experienced rats, 985 were classified as WS and 95 were classified as NS. Of the 1313 units recorded from the PL of cocaine-experienced rats, 1209 were classified as WS and 98 were classified as NS.

#### Neural firing rate dynamics

To analyze the general response properties of units, firing rates for each unit were computed in 50 ms bins, averaged across completed free- and forced-choice trials in all blocks, and peak normalized (Fig. 3a, c). Units were counted as maximally active for a trial epoch if, for at least one bin during that period, the unit’s peak-normalized average firing rate exceeded 95% of its absolute maximum value (Fig. 3b, d).

#### Individual unit regression analysis

Dynamics of neural selectivity for task variables were analyzed using linear regression models fit to each unit’s firing rate over the course of completed free- and forced-choice trials as previously described ^56; MATLAB^ ^function: fitglm;^ Fig. 4a, b. Within each epoch, firing rates were calculated in 50-ms bins, and for each bin, an individual regression model was computed. Outcome flavor, size, response direction and choice type served as categorical predictors. Within each unit, p-values were corrected for multiple comparisons, and the percent of the total population significant for each predictor was calculated for individual time bins (corrected p-value < 0.05).

#### Population and pseudoensemble analyses

Quantification of dimension representations – choice type, response direction, size and flavor – were calculated using the d’ metric in 100 unit pseudoensembles. Pseudoensembles were generated by randomly pulling subsets of 100 units for inclusion from the entire classified WS population per region of recorded units across all sessions. Pseudoensemble generation and d’ calculations were performed independently for sessions recorded from sucrose- and cocaine-experienced rats. Pseudoensembles were used to calculate median unit firing rates which were then z-scored across each of the 16 unique *choice type* x response *direction* x *size* x *flavor* combinations, creating population vectors for each combination. Correlations were then drawn between vectors associated with each trial-related dimension combination, and d’ metrics were computed measuring the separation of distributions of the correlations from within-dimension conditions (e.g., pairs of identical choice type combinations) versus the distribution of the correlations from between-dimension conditions (e.g., pairs of opposing choice type combinations). D’ values of each dimension were averaged across 250 runs, comparisons were made between dimension (choice type- vs. response direction- vs. size- vs. flavor), treatment groups (sucrose- vs. cocaine-experienced), and time (100 ms bins), and effects were analyzed using two-way ANOVAs (Fig. 5a, b).

To measure the degree to which multiple task dimensions were represented by cortical population ensembles during the odor sample epoch, WS units per region and treatment group were pooled and median unit firing rates were z-scored across each of the 16 unique *choice type* x response *direction* x *size* x *flavor* combinations, again creating population vectors for each combination. The relationship between ensemble representations were then measured by computing the correlation coefficient between population vectors for every pair of trial-related variable combinations ^57, 58^. Population level hierarchical representations were visualized using dendrograms (MATLAB functions: *linkage* and *dendrogram*) and cluster trees were computed using the unweighted average distance (UPGMA) between pairs of vectors and correlation as the distance metric (Fig. 6a, b).

To examine whether the strength of OFC WS choice representations during odor sample were related to across-choice-type transition errors, we first generated pseudoensembles by binning every two recording sessions for which each rat was run and the pooled unit population exceeded 20 units (n = 8 bins; i.e., first bin session: all rats’ sessions 2-3, second bin session: all rat’s sessions: 4-5, etc.) and pulled subsets of 20 classified OFC WS units per treatment group.

Population vectors were created using pseudoensembles for each of the eight bin sessions and d’ of choice type was calculated for each bin as described above using neural activity occurring during the odor sample epoch. Next, corresponding across-choice transition errors were calculated by averaging over the same sessions included in each bin. Finally, we fit a generalized linear model using d’ of choice type and treatment group as predictors and across-choice-type transition errors as the response to look at how well behavior could be explained by the strength (i.e., d’) of choice type representations (MATLAB function: fitglm; Fig. 6d).

The strength of latent representations were quantified using “shattering dimensionality” (SD, Fig. 7), a previously established population classification algorithm Fig. 7, ^59^. SD was performed per region and treatment group across all trial type combinations possible (i.e., 35 unique ways of evenly pairing eight trial types defined by response *direction (left, L or right, R) × size (small, S or big, B) × flavor* (chocolate, C or vanilla, V) trial mapping conditions). The three balanced dichotomies corresponding with the known task-related variables of response direction (i.e., LSC LBC LSV LBV vs. RSC RBC RSV RBV), size (i.e., LSC RSC LSV RSV vs LBC RBC LBV RBV), and flavor (i.e., LSC RSC RBC LBC vs. LSV RSV RBV LBV) were excluded, with the other 32 balanced arbitrary pairings (e.g., RSC LBC LSV LBV vs. LSC RBC RSV RBV) remaining as latent variables. WS units were included in the analysis if activity was recorded for at least 15 trials of each of the eight trial types on correctly-performed forced-choice trials, and spike counts were taken from a 450 ms window during the odor sample epoch. Testing and training vectors were created for each of the eight trial conditions possible, with 5 of the 15 randomly selected trials serving as testing sets, and the remaining trials serving as training sets. Associated training and testing spike counts were then normalized using means and standard deviations calculated from the training sets. To accomplish a number of trials greater than the number of units, normalized spike counts from training sets were randomly re-sampled for a total of 10,000 trials per trial type, creating single-trial population training vectors. Similarly, normalized spike counts from testing sets were randomly re-sampled for a total of 1,000 trials per trial type, creating single-trial population testing vectors. Binary SVM classifiers were then trained and tested using the single-trial population vectors to compute how well trial type could be decoded from activity. The entire procedure was repeated 1000 times and then averaged across all repetitions for each of the 32 balanced arbitrary pairings.

#### Statistical Analyses

All analyses were conducted using MATLAB (The MathWorks Inc., Natick, MA) and GraphPad Prism 8 software (GraphPad, La Jolla, CA). One- and two-way ANOVAs, ANCOVAs, and t-tests were used to analyze all data, as reported in results and figure legends. The Bonferroni procedure was used to correct for multiple comparisons. P-values were significant if they fell below 0.05. Sample sizes were determined from previous studies utilizing similar behavioral and recording procedures ^42, 44, 56, 84^. Error bars or shading on plots represent the standard error of the mean.

## Figures and legends

**Supplemental Figure 1.**
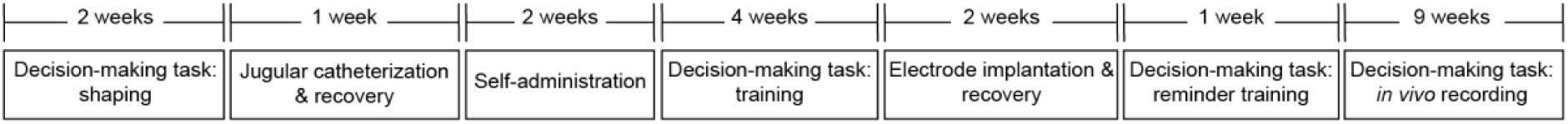
Experimental timeline.

**Supplemental Figure 2.**
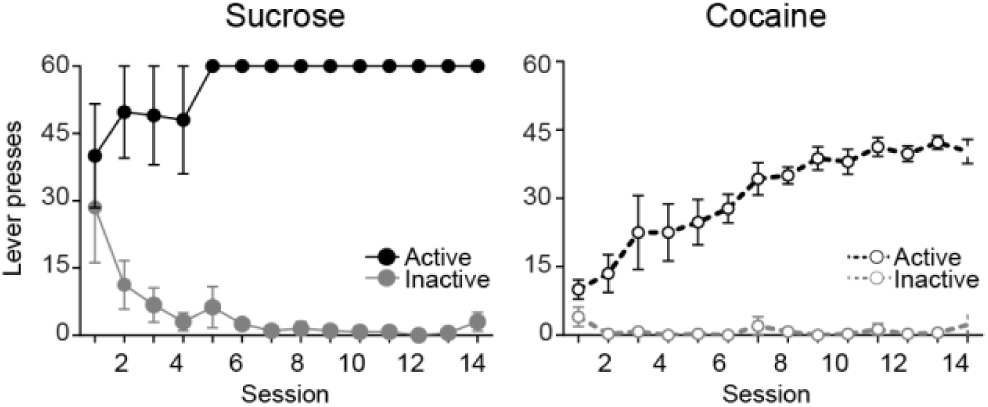
Rats were trained to self-administer either oral sucrose (n = 4) or cocaine (n = 4) using a fixed ratio one schedule of reinforcement for 3 h/d for 14 d. Plotted are the number of active (black circles) and inactive (grey circles) lever presses during each session of sucrose (left; n = 4) and cocaine (right; n = 4) self-administration. A two-way repeated-measures ANOVA detected a significant session × lever interaction in both treatment groups: sucrose, *P* < 0.0001, F(13,39) = 7.348; cocaine, *P* < 0.0001, F(13,39) = 20.21), suggesting that both groups acquired the task. Error bars indicate SEM in lever presses.

**Supplemental Figure 3.**
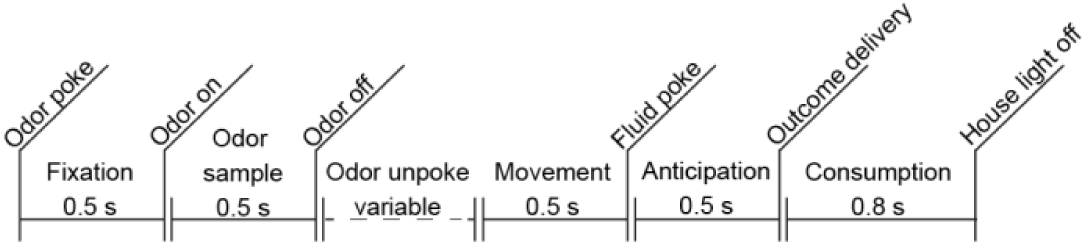
Completed free- and forced-choice trials were divided into five epochs bounded by important task events.

## References

1. APA. Diagnostic and statistical manual of mental disorders (Arlington, VA, 2013).

2. Koffarnus, M.N. & Kaplan, B.A. Clinical models of decision making in addiction. Pharmacol Biochem Behav 164, 71–83 (2018).

3. SAMHSA. Mental and Substance Use Disorders. (2016).

4. Fillmore, M.T. & Rush, C.R. Polydrug abusers display impaired discrimination-reversal learning in a model of behavioural control. J Psychopharmacol 20, 24–32 (2006).

5. Verdejo-Garcia, A.J., Perales, J.C. & Perez-Garcia, M. Cognitive impulsivity in cocaine and heroin polysubstance abusers. Addict Behav 32, 950–966 (2007).

6. Calu, D.J., et al. Withdrawal from cocaine self-administration produces long-lasting deficits in orbitofrontal-dependent reversal learning in rats. Learn Mem 14, 325–328 (2007).

7. Schoenbaum, G., Saddoris, M.P., Ramus, S.J., Shaham, Y. & Setlow, B. Cocaine-experienced rats exhibit learning deficits in a task sensitive to orbitofrontal cortex lesions. Eur J Neurosci 19, 1997–2002 (2004).

8. Krueger, D.D., et al. Prior chronic cocaine exposure in mice induces persistent alterations in cognitive function. Behav Pharmacol 20, 695–704 (2009).

9. Jentsch, J.D., Olausson, P., De La Garza, R., 2nd & Taylor, J.R. Impairments of reversal learning and response perseveration after repeated, intermittent cocaine administrations to monkeys. Neuropsychopharmacology 26, 183–190 (2002).

10. Porter, J.N., et al. Chronic cocaine self-administration in rhesus monkeys: impact on associative learning, cognitive control, and working memory. J Neurosci 31, 4926–4934 (2011).

11. Nelson, A. & Killcross, S. Amphetamine exposure enhances habit formation. J Neurosci 26, 3805–3812 (2006).

12. Schoenbaum, G. & Setlow, B. Cocaine makes actions insensitive to outcomes but not extinction: implications for altered orbitofrontal-amygdalar function. Cereb Cortex 15, 1162–1169 (2005).

13. Dickinson, A., Wood, N. & Smith, J.W. Alcohol seeking by rats: action or habit? Q J Exp Psychol B 55, 331–348 (2002).

14. Miles, F.J., Everitt, B.J. & Dickinson, A. Oral cocaine seeking by rats: action or habit? Behav Neurosci 117, 927–938 (2003).

15. Wied, H.M., Jones, J.L., Cooch, N.K., Berg, B.A. & Schoenbaum, G. Disruption of model-based behavior and learning by cocaine self-administration in rats. Psychopharmacology (Berl) 229, 493–501 (2013).

16. Marshall, A.T. & Ostlund, S.B. Repeated cocaine exposure dysregulates cognitive control over cue-evoked reward-seeking behavior during Pavlovian-to-instrumental transfer. Learn Mem 25, 399–409 (2018).

17. Izquierdo, A., Suda, R.K. & Murray, E.A. Bilateral orbital prefrontal cortex lesions in rhesus monkeys disrupt choices guided by both reward value and reward contingency. J Neurosci 24, 7540–7548 (2004).

18. Gallagher, M., McMahan, R.W. & Schoenbaum, G. Orbitofrontal cortex and representation of incentive value in associative learning. J Neurosci 19, 6610–6614 (1999).

19. Shipman, M.L., Trask, S., Bouton, M.E. & Green, J.T. Inactivation of prelimbic and infralimbic cortex respectively affects minimally-trained and extensively-trained goal-directed actions. Neurobiol Learn Mem 155, 164–172 (2018).

20. Jones, J.L., et al. Orbitofrontal cortex supports behavior and learning using inferred but not cached values. Science 338, 953–956 (2012).

21. Li, J.S., Hsiao, K.Y. & Chen, W.M. Effects of medial prefrontal cortex lesions in rats on the what-where-when memory of a fear conditioning event. Behav Brain Res 218, 94–98 (2011).

22. Takahashi, Y.K., et al. The orbitofrontal cortex and ventral tegmental area are necessary for learning from unexpected outcomes. Neuron 62, 269–280 (2009).

23. Burke, K.A., Takahashi, Y.K., Correll, J., Brown, P.L. & Schoenbaum, G. Orbitofrontal inactivation impairs reversal of Pavlovian learning by interfering with ‘disinhibition’ of responding for previously unrewarded cues. Eur J Neurosci 30, 1941–1946 (2009).

24. Naneix, F., Marchand, A.R., Di Scala, G., Pape, J.R. & Coutureau, E. A role for medial prefrontal dopaminergic innervation in instrumental conditioning. J Neurosci 29, 6599–6606 (2009).

25. Balleine, B.W. & Dickinson, A. Goal-directed instrumental action: contingency and incentive learning and their cortical substrates. Neuropharmacology 37, 407–419 (1998).

26. Corbit, L.H. & Balleine, B.W. The role of prelimbic cortex in instrumental conditioning. Behav Brain Res 146, 145–157 (2003).

27. Jackson, S.A., Horst, N.K., Pears, A., Robbins, T.W. & Roberts, A.C. Role of the Perigenual Anterior Cingulate and Orbitofrontal Cortex in Contingency Learning in the Marmoset. Cereb Cortex 26, 3273–3284 (2016).

28. Schiller, D. & Weiner, I. Lesions to the basolateral amygdala and the orbitofrontal cortex but not to the medial prefrontal cortex produce an abnormally persistent latent inhibition in rats. Neuroscience 128, 15–25 (2004).

29. Costa, K.M., Sengupta, A. & Schoenbaum, G. The orbitofrontal cortex is necessary for learning to ignore. Curr Biol 31, 2652–2657.e2653 (2021).

30. George, D.N., Duffaud, A.M., Pothuizen, H.H., Haddon, J.E. & Killcross, S. Lesions to the ventral, but not the dorsal, medial prefrontal cortex enhance latent inhibition. Eur J Neurosci 31, 1474–1482 (2010).

31. Ostlund, S.B. & Balleine, B.W. Orbitofrontal cortex mediates outcome encoding in Pavlovian but not instrumental conditioning. J Neurosci 27, 4819–4825 (2007).

32. Homayoun, H. & Moghaddam, B. Differential representation of Pavlovian-instrumental transfer by prefrontal cortex subregions and striatum. Eur J Neurosci 29, 1461–1476 (2009).

33. Halbout, B., Hutson, C., Wassum, K.M. & Ostlund, S.B. Dorsomedial prefrontal cortex activation disrupts Pavlovian incentive motivation. Front Behav Neurosci 16, 999320 (2022).

34. Gobin, C., Shallcross, J. & Schwendt, M. Neurobiological substrates of persistent working memory deficits and cocaine-seeking in the prelimbic cortex of rats with a history of extended access to cocaine self-administration. Neurobiol Learn Mem 161, 92–105 (2019).

35. Jentsch, J.D. & Taylor, J.R. Impulsivity resulting from frontostriatal dysfunction in drug abuse: implications for the control of behavior by reward-related stimuli. Psychopharmacology (Berl) 146, 373–390 (1999).

36. Volkow, N.D. & Fowler, J.S. Addiction, a disease of compulsion and drive: involvement of the orbitofrontal cortex. Cereb Cortex 10, 318–325 (2000).

37. Bechara, A., et al. Decision-making deficits, linked to a dysfunctional ventromedial prefrontal cortex, revealed in alcohol and stimulant abusers. Neuropsychologia 39, 376–389 (2001).

38. Ersche, K.D., Roiser, J.P., Robbins, T.W. & Sahakian, B.J. Chronic cocaine but not chronic amphetamine use is associated with perseverative responding in humans. Psychopharmacology (Berl) 197, 421–431 (2008).

39. Grant, S., Contoreggi, C. & London, E.D. Drug abusers show impaired performance in a laboratory test of decision making. Neuropsychologia 38, 1180–1187 (2000).

40. Chen, B.T., et al. Rescuing cocaine-induced prefrontal cortex hypoactivity prevents compulsive cocaine seeking. Nature 496, 359–362 (2013).

41. Homayoun, H. & Moghaddam, B. Progression of cellular adaptations in medial prefrontal and orbitofrontal cortex in response to repeated amphetamine. J Neurosci 26, 8025–8039 (2006).

42. Stalnaker, T.A., Roesch, M.R., Franz, T.M., Burke, K.A. & Schoenbaum, G. Abnormal associative encoding in orbitofrontal neurons in cocaine-experienced rats during decision-making. Eur J Neurosci 24, 2643–2653 (2006).

43. Thorpe, S.J., Rolls, E.T. & Maddison, S. The orbitofrontal cortex: neuronal activity in the behaving monkey. Exp Brain Res 49, 93–115 (1983).

44. Stalnaker, T.A., Berg, B., Aujla, N. & Schoenbaum, G. Cholinergic Interneurons Use Orbitofrontal Input to Track Beliefs about Current State. J Neurosci 36, 6242–6257 (2016).

45. Letzkus, J.J., et al. A disinhibitory microcircuit for associative fear learning in the auditory cortex. Nature 480, 331–335 (2011).

46. Quirk, M.C., Sosulski, D.L., Feierstein, C.E., Uchida, N. & Mainen, Z.F. A defined network of fast-spiking interneurons in orbitofrontal cortex: responses to behavioral contingencies and ketamine administration. Front Syst Neurosci 3, 13 (2009).

47. Sotres-Bayon, F., Sierra-Mercado, D., Pardilla-Delgado, E. & Quirk, G.J. Gating of fear in prelimbic cortex by hippocampal and amygdala inputs. Neuron 76, 804–812 (2012).

48. Insel, N. & Barnes, C.A. Differential Activation of Fast-Spiking and Regular-Firing Neuron Populations During Movement and Reward in the Dorsal Medial Frontal Cortex. Cereb Cortex 25, 2631–2647 (2015).

49. Cauli, B., et al. Classification of fusiform neocortical interneurons based on unsupervised clustering. Proc Natl Acad Sci U S A 97, 6144–6149 (2000).

50. Kawaguchi, Y. & Kubota, Y. GABAergic cell subtypes and their synaptic connections in rat frontal cortex. Cereb Cortex 7, 476–486 (1997).

51. McCormick, D.A., Connors, B.W., Lighthall, J.W. & Prince, D.A. Comparative electrophysiology of pyramidal and sparsely spiny stellate neurons of the neocortex. J Neurophysiol 54, 782–806 (1985).

52. Feierstein, C.E., Quirk, M.C., Uchida, N., Sosulski, D.L. & Mainen, Z.F. Representation of spatial goals in rat orbitofrontal cortex. Neuron 51, 495–507 (2006).

53. Roesch, M.R., Taylor, A.R. & Schoenbaum, G. Encoding of time-discounted rewards in orbitofrontal cortex is independent of value representation. Neuron 51, 509–520 (2006).

54. Stalnaker, T.A., et al. Orbitofrontal neurons infer the value and identity of predicted outcomes. Nat Commun 5, 3926 (2014).

55. Tsujimoto, S., Genovesio, A. & Wise, S.P. Monkey orbitofrontal cortex encodes response choices near feedback time. J Neurosci 29, 2569–2574 (2009).

56. Wikenheiser, A.M., Marrero-Garcia, Y. & Schoenbaum, G. Suppression of Ventral Hippocampal Output Impairs Integrated Orbitofrontal Encoding of Task Structure. Neuron 95, 1197–1207.e1193 (2017).

57. McKenzie, S., et al. Hippocampal representation of related and opposing memories develop within distinct, hierarchically organized neural schemas. Neuron 83, 202–215 (2014).

58. Kriegeskorte, N., Mur, M. & Bandettini, P. Representational similarity analysis - connecting the branches of systems neuroscience. Front Syst Neurosci 2, 4 (2008).

59. Bernardi, S., et al. The Geometry of Abstraction in the Hippocampus and Prefrontal Cortex. Cell 183, 954–967.e921 (2020).

60. Schoenbaum, G., Setlow, B., Nugent, S.L., Saddoris, M.P. & Gallagher, M. Lesions of orbitofrontal cortex and basolateral amygdala complex disrupt acquisition of odor-guided discriminations and reversals. Learn Mem 10, 129–140 (2003).

61. Schoenbaum, G., Chiba, A.A. & Gallagher, M. Neural encoding in orbitofrontal cortex and basolateral amygdala during olfactory discrimination learning. J Neurosci 19, 1876–1884 (1999).

62. Wallis, J.D. & Miller, E.K. Neuronal activity in primate dorsolateral and orbital prefrontal cortex during performance of a reward preference task. Eur J Neurosci 18, 2069–2081 (2003).

63. Birrell, J.M. & Brown, V.J. Medial frontal cortex mediates perceptual attentional set shifting in the rat. J Neurosci 20, 4320–4324 (2000).

64. Marquis, J.P., Killcross, S. & Haddon, J.E. Inactivation of the prelimbic, but not infralimbic, prefrontal cortex impairs the contextual control of response conflict in rats. Eur J Neurosci 25, 559–566 (2007).

65. Floresco, S.B., Block, A.E. & Tse, M.T. Inactivation of the medial prefrontal cortex of the rat impairs strategy set-shifting, but not reversal learning, using a novel, automated procedure. Behav Brain Res 190, 85–96 (2008).

66. Ragozzino, M.E., Kim, J., Hassert, D., Minniti, N. & Kiang, C. The contribution of the rat prelimbic-infralimbic areas to different forms of task switching. Behav Neurosci 117, 1054–1065 (2003).

67. Stefani, M.R., Groth, K. & Moghaddam, B. Glutamate receptors in the rat medial prefrontal cortex regulate set-shifting ability. Behav Neurosci 117, 728–737 (2003).

68. Stefani, M.R. & Moghaddam, B. Systemic and prefrontal cortical NMDA receptor blockade differentially affect discrimination learning and set-shift ability in rats. Behav Neurosci 119, 420–428 (2005).

69. Schoenbaum, G. & Shaham, Y. The role of orbitofrontal cortex in drug addiction: a review of preclinical studies. Biol Psychiatry 63, 256–262 (2008).

70. Everitt, B.J., et al. The orbital prefrontal cortex and drug addiction in laboratory animals and humans. Ann N Y Acad Sci 1121, 576–597 (2007).

71. Lucantonio, F., et al. Orbitofrontal activation restores insight lost after cocaine use. Nat Neurosci 17, 1092–1099 (2014).

72. Panayi, M.C., et al. The selective D3-Receptor antagonist VK4-116 effectively treats behavioral inflexibility in rats caused by self-administration and withdrawal from cocaine. bioRxiv (2023).

73. Sadacca, B.F., et al. Orbitofrontal neurons signal sensory associations underlying model-based inference in a sensory preconditioning task. Elife 7 (2018).

74. Hogarth, L., et al. Intact goal-directed control in treatment-seeking drug users indexed by outcome-devaluation and Pavlovian to instrumental transfer: critique of habit theory. Eur J Neurosci 50, 2513–2525 (2019).

75. Song, M., et al. Minimal cross-trial generalization in learning the representation of an odor-guided choice task. PLoS Comput Biol 18, e1009897 (2022).

76. Baltz, E.T., Yalcinbas, E.A., Renteria, R. & Gremel, C.M. Orbital frontal cortex updates state-induced value change for decision-making. Elife 7 (2018).

77. Bradfield, L.A., Bertran-Gonzalez, J., Chieng, B. & Balleine, B.W. The thalamostriatal pathway and cholinergic control of goal-directed action: interlacing new with existing learning in the striatum. Neuron 79, 153–166 (2013).

78. Schuck, N.W., Cai, M.B., Wilson, R.C. & Niv, Y. Human Orbitofrontal Cortex Represents a Cognitive Map of State Space. Neuron 91, 1402–1412 (2016).

79. Wilson, R.C., Takahashi, Y.K., Schoenbaum, G. & Niv, Y. Orbitofrontal cortex as a cognitive map of task space. Neuron 81, 267–279 (2014).

80. Gardner, M.P.H. & Schoenbaum, G. The orbitofrontal cartographer. Behav Neurosci 135, 267–276 (2021).

81. Costa, K.M., et al. The role of the lateral orbitofrontal cortex in creating cognitive maps. Nat Neurosci 26, 107–115 (2023).

82. Stalnaker, T.A., Calhoon, G.G., Ogawa, M., Roesch, M.R. & Schoenbaum, G. Neural correlates of stimulus-response and response-outcome associations in dorsolateral versus dorsomedial striatum. Front Integr Neurosci 4, 12 (2010).

83. Burgos-Robles, A., Bravo-Rivera, H. & Quirk, G.J. Prelimbic and infralimbic neurons signal distinct aspects of appetitive instrumental behavior. PloS one 8, e57575–e57575 (2013).

84. Mueller, L.E., Sharpe, M.J., Stalnaker, T.A., Wikenheiser, A.M. & Schoenbaum, G. Prior Cocaine Use Alters the Normal Evolution of Information Coding in Striatal Ensembles during Value-Guided Decision-Making. J Neurosci 41, 342–353 (2021).

